# Limits on Inferring Gene Regulatory Networks Subjected to Different Noise Mechanisms

**DOI:** 10.1101/2023.01.23.525259

**Authors:** Michael Saint-Antoine, Abhyudai Singh

## Abstract

One of the most difficult and pressing problems in computational cell biology is the inference of gene regulatory network structure from transcriptomic data. Benchmarking network inference methods on model organism datasets has yielded mixed results, in which the methods sometimes perform reasonably well and other times fail to outperform random guessing. In this paper, we analyze the feasibility of network inference under different noise conditions using stochastic simulations. We show that gene regulatory interactions with extrinsic noise appear to be more amenable to inference than those with only intrinsic noise, especially when the extrinsic noise causes the system to switch between distinct expression states. Furthermore, we analyze the problem of false positives between genes that have no direct interaction but share a common upstream regulator, and explore a strategy for distinguishing between these false positives and true interactions based on noise profiles of mRNA expression levels. Lastly, we derive mathematical formulas for the mRNA noise levels and correlation using moment analysis techniques, and show how these levels change as the mean mRNA expression level changes.

## I. Introduction

In what is known as the “central dogma” of molecular biology, information encoded in DNA is transcribed into strands of messenger RNA (mRNA), which are then translated into proteins, which then carry out various functions within the cell. Sometimes the function of a protein from one gene involves regulating the expression of other genes. When a protein increases the expression of another gene, we refer to this as “activation.” When a protein decreases the expression of another gene, we refer to this as “repression.” Intricate networks of these positive and negative regulatory interactions between genes (called gene regulatory networks or GRNs) give rise to much of the complexity of life [17], [27]. Understanding the roles and functionality of GRNs is a pressing problem for cell biologists, especially since malfunctions of GRNs can have disastrous medical impacts on human health, leading to diseases like cancer, for example [19].

In this paper, we present models of different noise conditions related to regulatory interactions between genes. In this context, *intrinsic noise* refers to the inherent stochasticity in the processes of transcription, translation, and the degradation of mRNA and proteins, and is especially prevalent in cases of low copy number fluctuations of these molecules. By contrast, *extrinsic noise* refers to the impact of other factors, such as upstream regulators, external stimuli, or changes in cell state that affect the regulatory interaction [6], [13], [32], [36], [38].

A specific challenge for computational biologists studying GRNs is network inference – that is, the attempt to infer the structure of a GRN from gene expression data [29]. Although modern high-throughput next generation sequencing (NGS) experiments like RNA-seq have led to an abundance of gene expression data, the challenge of network inference is still quite difficult. Part of the reason for this difficulty is that NGS transcriptomic experiments like RNA-seq involve destroying each cell to sequence its RNA content. So, each cell provides only a single time point of data, rather than a timeseries dataset.

Most network inference methods attempt to infer regulatory interactions between genes based on statistical relationships between their expression levels (typically quantified by mRNA abundance). Some examples of these methods include correlation [45], linear or non-linear regression [11], [14], [37], information theory [4], [5], [7], [24], [25], Bayesian techniques [8], [44], and others [1]–[3], [15], [18], [43], [46].

An excellent introductory review of the topic of gene regulatory network inference can be found in Huynh-Thu and Sanguinetti 2018 [16]. Recent theoretical work on models of gene expression and regulation can be found in [22], [39]–[41].

## II. Efficacy of Network Inference Methods

Although the problem of gene regulatory network inference has been widely studied for more than a decade, there are still questions about the efficacy of these methods and whether network inference from transcriptomic data is a feasible goal. A key point of skepticism is that these methods typically assume that mRNA abundance measurements can be used as a reliable proxy for protein abundance. Typically, the protein (not the mRNA) produced by a gene is what regulates the expression of other genes, but it is often mRNA abundance data that we have access to, so most GRN inference methods take mRNA data as an input.

There are some reasons to question the assumption that mRNA abundance data can be used as a reliable proxy for protein abundance. For example, Mahajan et al. 2022 [21] shows though theoretical analysis and stochastic simulations that, under conditions of only intrinsic noise, the correlation between mRNA abundance and protein abundance *even for the same gene* becomes quite weak if there is a large difference between the mRNA stability and protein stability. Additionally, Liu et al. 2016 [20] reviews the literature and reports a similar finding, that the correlation between mRNA levels and protein levels can be weak in some scenarios, and knowledge of mRNA transcript abundance alone is not always sufficient to predict protein abundance levels.

So how well do these network inference methods actually work? There have been several attempts to test the efficacy of network inference methods by benchmarking them on data from model organisms, such as *E. coli*, *S. cerevisiae*, and mice [23], [26], [28]. In these benchmarking studies, the underlying structure of the gene regulatory network is already known from experimental investigation, so the predictions of network inference methods can be checked against the correct answers. The most famous of these benchmarking attempts is Marbach et al. 2012 [23]. The results of this benchmarking study were mixed. When tested on a *S. cerevisiae* dataset, network predictions failed to substantially outperform the accuracy that would be expected by random guessing. However, when tested on *E. coli* data, the network predictions performed substantially better than random guessing.

So, our current understanding of the efficacy of gene regulatory network inference is quite murky and uncertain. It seems that network inference from transcriptomic data cannot be considered entirely feasible or unfeasible. Rather, it seems to be feasible under some conditions and unfeasible under other conditions. In this paper, we attempt to shed light on this topic, investigating through stochastic simulations which noise conditions may be more or less amenable to network inference from transcriptomic data.

## III. Standard Activation Model, no Extrinsic Noise

In this section, we briefly review and replicate results from Mahajan et al. 2022 [21], which studied the feasibility of network inference from mRNA abundance data under conditions of only intrinsic noise. We consider a system with two genes: Gene 1 and Gene 2. Gene 1 is transcribed into mRNA1, which is then translated into the Protein. The Protein then activates the transcription of Gene 2 into mRNA2. We refer to this as the Activation scenario. A diagram of this scenario is shown in Figure 1.

**Fig. 1.**
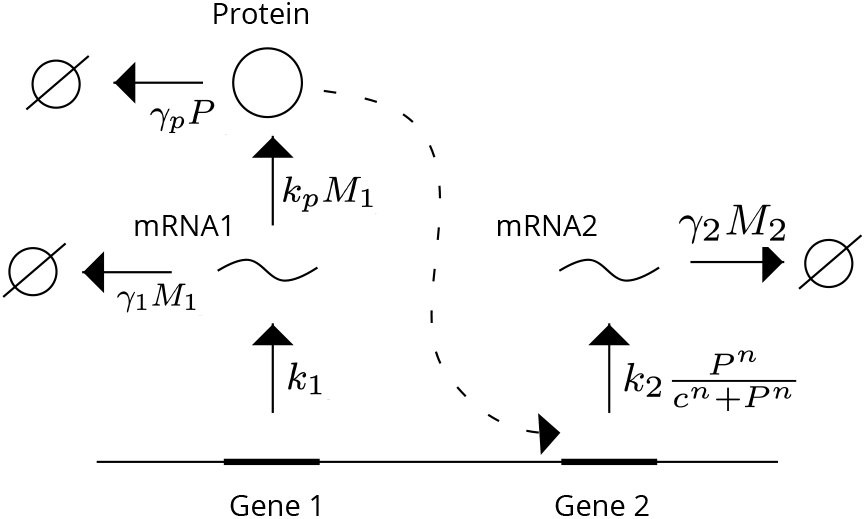
Activation Model, No Extrinsic Noise. *M*_1_ tracks the count of mRNA1, which is produced with a constant rate, and also degrades with some rate. *P* tracks the count of the Protein, which is produced from mRNA1, and alsoPrdoetgeriandes with some rate. The Protein activates the production of mRNA2, which also degrades with some rate. We track the count of mRNA2 with the variable *M*_2_.

We define the integer-valued random processes *M*_1_(*t*), *P* (*t*), and *M*_2_(*t*) to track the counts of the mRNA1, Protein, and mRNA2 respectively. For the sake of simplicity, we will refer to these processes as *M*_1_, *P*, and *M*_2_ from now on.

The stochastic model is described Table I. The model consists of six events that occur probabilistically with rates given in the third column. When the event occurs, the counts for the variables are updated according to the reset map in the second column. Descriptions of the parameters, as well as the values we used in our simulations, are listed in Table II. With this setup, we can simulate the model using Gillespie’s stochastic simulation algorithm (SSA) [9].

**TABLE I.**
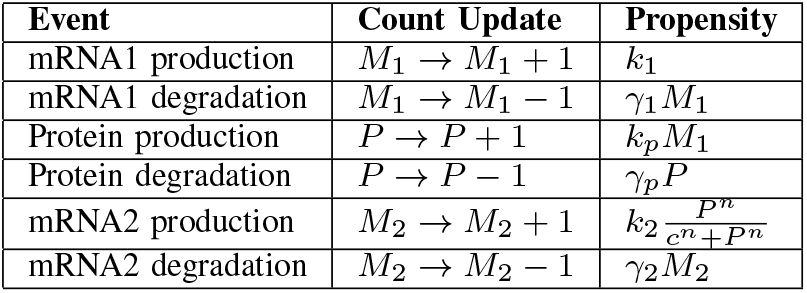
Activation Model, No Extrinsic Noise

**TABLE II.** Activation Model, Parameters

| Parameter | Description | Value |
| --- | --- | --- |
| $k_1$ | mRNA1 production rate | 4 |
| $\gamma_1$ | mRNA1 degradation rate | 2 |
| $k_p$ | Protein production rate | 2 |
| $\gamma_p$ | Protein degradation rate | $\frac{2}{\tau}$ |
| $k_2$ | mRNA2 maximum production rate | 4 |
| $\gamma_2$ | mRNA2 degradation rate | 2 |
| $n$ | Hill coefficient | 3 |
| $c$ | Hill function parameter | 1 |

We model the activation of *M*_2_ production by *P* as the Hill function 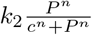, where *P* is the level of the Protein, *n* is the Hill coefficient (which determines how linear or nonlinear the activation is), *c* is a constant parameter that affects the saturation dynamics of the Hill function, and *k*_2_ is the maximum production rate. As *P* increases, the fraction 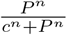 saturates and approaches 1, so the entire term 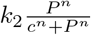 approaches *k*_2_

Part of the analysis in [21] involved calculating the correlation between the mRNA levels under different assumptions about the relative stability of the mRNA and protein. In our case, we are intereGsetneed1in the correlaGtieonne 2between *M*_1_ and *M*_2_ under different assumptions about the relative stability of mRENxtArin1sicand the Protein. The stability of mRNA1 is the reciproFaccatlorof its degradation rate: 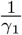 The stability of the Protein, likewise, is: 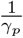. So, the ratio of Protein stability to mRNA1 stability can be expressed as 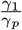. We refer to this stability ratio as *τ*.

A key finding in [21] was that in this model with only intrinsic noise, correlation between the mRNA levels is quite weak, and gets weaker as the ratio of stability between the protein and mRNA increases. Figure 2 shows our replication of this result: the correlation between *M*_1_ and *M*_2_ is quite weak, and drops to nearly 0 as the ratio of stability between the Protein and mRNA1 increases.

**Fig. 2.**
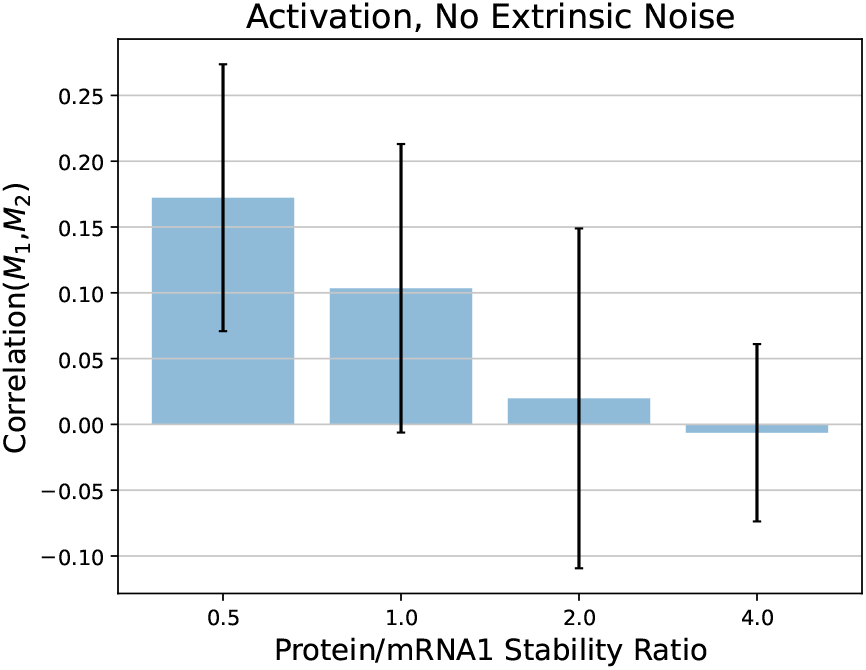
Plot of the Pearson correlation coefficient between *M*_1_ and *M*_2_ for different ratios of Protein stability to mRNA1 stability, which we define in terms of the degradation rates as 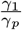. Error bars show one standard deviation. Simulations were run using the parameter values shown in Table II.

This result seems to give a bleak outlook for the challenge of network inference. If there is weak or zero correlation between mRNA abundance for genes that regulate each other, how can we hope to infer gene regulatory network structure from transcriptomic data? However, as noted in the previous section, attempts to benchmark network inference methods on real data have yielded mixed results. In some cases, the network inference methods sometimes have performed reasonably well, and in other cases they have failed to outperform random guessing. In the next section, we will modify the model in a way that could explain this mixed-feasibility of network inference.

## IV. Cascade Model With Extrinsic Noise

In the previous section, we analyzed a model that included only intrinsic noise in the processes of transcription and translation. However, a more realistic model of the biological system might include extrinsic noise, which could come from environmental stimuli, changes to the internal cell state, or regulation from another upstream gene. In this section, we introduce a new component to the model, which we refer to as the Extrinsic Factor, to represent the extrinsic noise source. We track the level of the Extrinsic Factor with the integer-valued random process *Z*(*t*). From now on we will refer to *Z*(*t*) as simply *Z* for the sake of simplicity.

In this model, Gene 1 is transcribed mRNA1 (with counts tracked by *M*_1_) which is then translated into the Protein (with counts tracked by *P*), which then activates the transcription of Gene 2 into mRNA2 (with counts tracked by *M*_2_). However, unlike the model in the previous section, in this model the production mRNA1 is activated by the Extrinsic Factor. We purposely define the Extrinsic Factor in vague biological terms, so that it can be thought of as an upstream transcription factor, external stimulus, or any other source of extrinsic noise affecting the transcription of mRNA1. We refer to this as the Cascade scenario. A diagram of this scenario is shown in Figure 3.

**Fig. 3.**
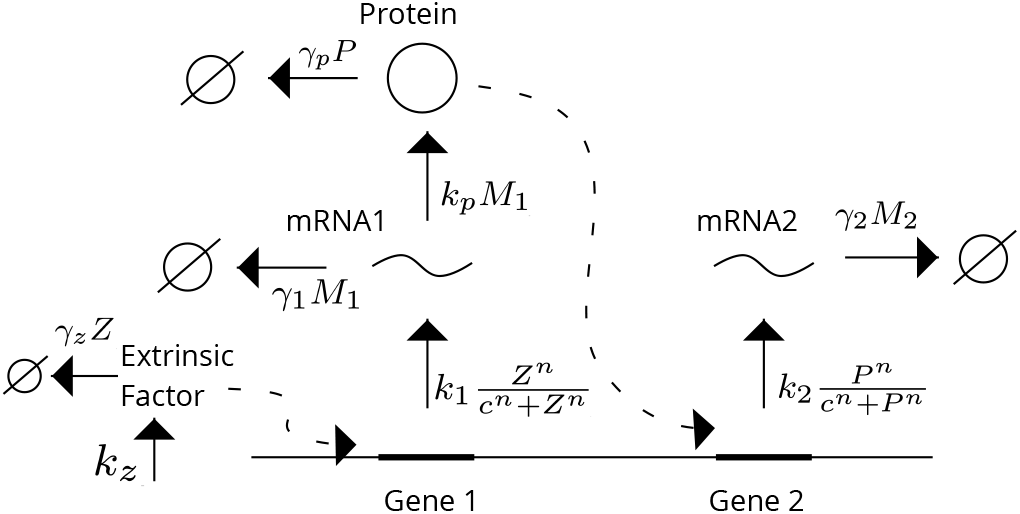
Cascade Model with Extrinsic Noise. We introduce an Extrinsic Factor (tracked by *Z*) to the model, which we purposely define in abstract terms so that it can refer to an upstream transcription factor, environmental stimulus, or something else. The Extrinsic Factor activates the production of mRNA1 (tracked by *M*_1_), which also degrades with some rate. The Protein (tracked by *P*) is produced from mRNA1, and also degrades with some rate. The Protein activates the production of mRNA2 (tracked by *M*_2_), which also degrades with some rate.

This stochastic model is described in Table III. Unlike the other variables, which update with increases and decreases of 1, the production of the Extrinsic Factor occurs with a burst of size *β*, which is drawn from a random geometric distribution with mean 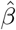. With this setup, increasing 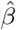 while holding the mean of *Z* constant increases the noise level of *Z*. In this section, we will report results for different mean burst sizes of 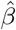, ranging 1 to 20. In all of these cases, the mean burst size 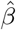 is changed, and the parameter *k_z_* is updated so that the mean of *Z* over time is held constant. We model the degradation of the Extrinsic Factor with the parameter *γ_z_*. In this model, the production of mRNA1 is activated by the Extrinsic Factor via a Hill function, with a maximum production limit of *κ*_1_. The Hill function parameters *c* and *n* are the same for both the activation mRNA1 transcription and the activation of mRNA2 transcription. Other than these changes, we model the degradation of mRNA1, the production and degradation of the Protein, and the production and degradation of mRNA2 the same as in the previous section.

**TABLE III.**
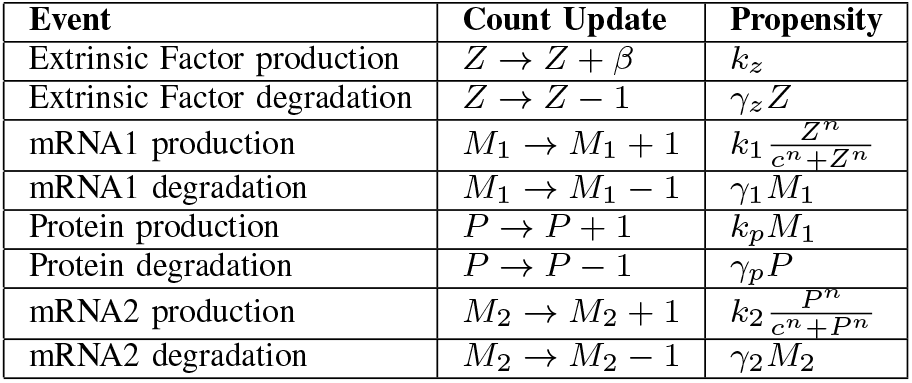
Cascade Model With Extrinsic Noise

| Event | Count Update | Propensity |
| --- | --- | --- |
| Extrinsic Factor production | $Z \rightarrow Z + \beta$ | $k_z$ |
| Extrinsic Factor degradation | $Z \rightarrow Z - 1$ | $\gamma_z Z$ |
| mRNA1 production | $M_1 \rightarrow M_1 + 1$ | $k_1 \frac{Z^n}{c^n + Z^n}$ |
| mRNA1 degradation | $M_1 \rightarrow M_1 - 1$ | $\gamma_1 M_1$ |
| Protein production | $P \rightarrow P + 1$ | $k_p M_1$ |
| Protein degradation | $P \rightarrow P - 1$ | $\gamma_p P$ |
| mRNA2 production | $M_2 \rightarrow M_2 + 1$ | $k_2 \frac{P^n}{c^n + P^n}$ |
| mRNA2 degradation | $M_2 \rightarrow M_2 - 1$ | $\gamma_2 M_2$ |

Figures 4, 5, and 6 show three scenarios in which the mean of *Z* is held constant, but the burst size mean 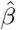 is varied. In Figure 4, the burst size is only 1, and in this simulation the correlation between *M*_1_ and *M*_2_ is 0.109. In Figure 5, the mean burst size is 10, and in this simulation the correlation between *M*_1_ and *M*_2_ is 0.334. In Figure 6, the mean burst size is 20, and in this simulation the correlation between *M*_1_ and *M*_2_ is 0.469.

**Fig. 4.**
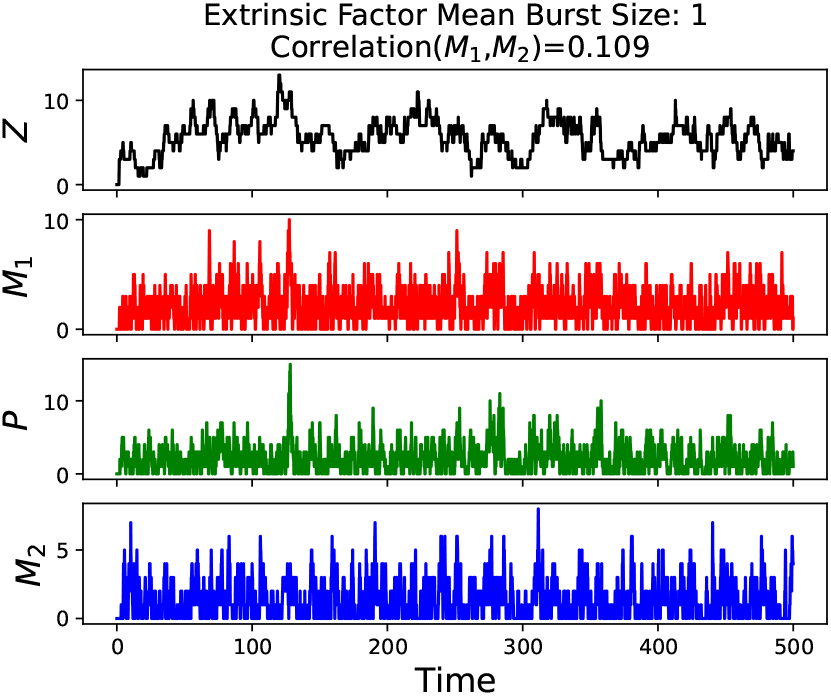
Simulation using parameter values listed in Table IV, with mean 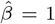, meaning that *β* = 1 with no variation. This simulation yielded a correlation coefficient between *M*_1_ and *M*_2_ of 0.109.

**TABLE IV.**
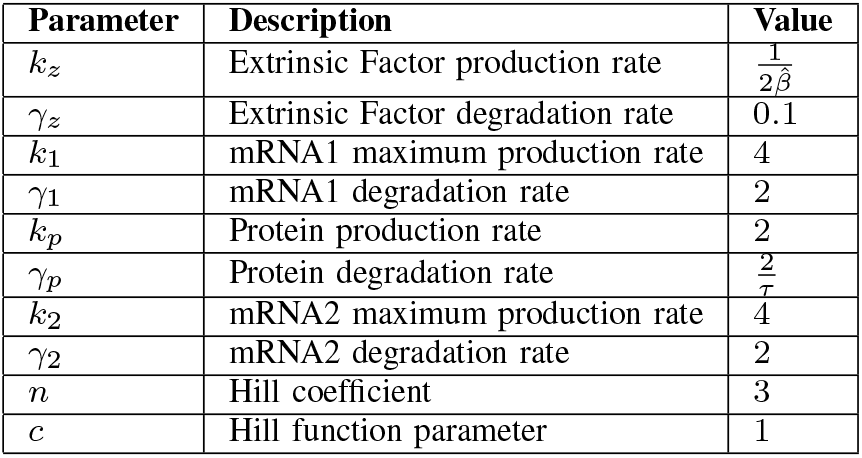
Cascade Model Parameters

| Parameter | Description | Value |
| --- | --- | --- |
| $k_z$ | Extrinsic Factor production rate | $\frac{1}{2\hat{\beta}}$ |
| $\gamma_z$ | Extrinsic Factor degradation rate | 0.1 |
| $k_1$ | mRNA1 maximum production rate | 4 |
| $\gamma_1$ | mRNA1 degradation rate | 2 |
| $k_p$ | Protein production rate | 2 |
| $\gamma_p$ | Protein degradation rate | $\frac{2}{\tau}$ |
| $k_2$ | mRNA2 maximum production rate | 4 |
| $\gamma_2$ | mRNA2 degradation rate | 2 |
| $n$ | Hill coefficient | 3 |
| $c$ | Hill function parameter | 1 |

**Fig. 5.**
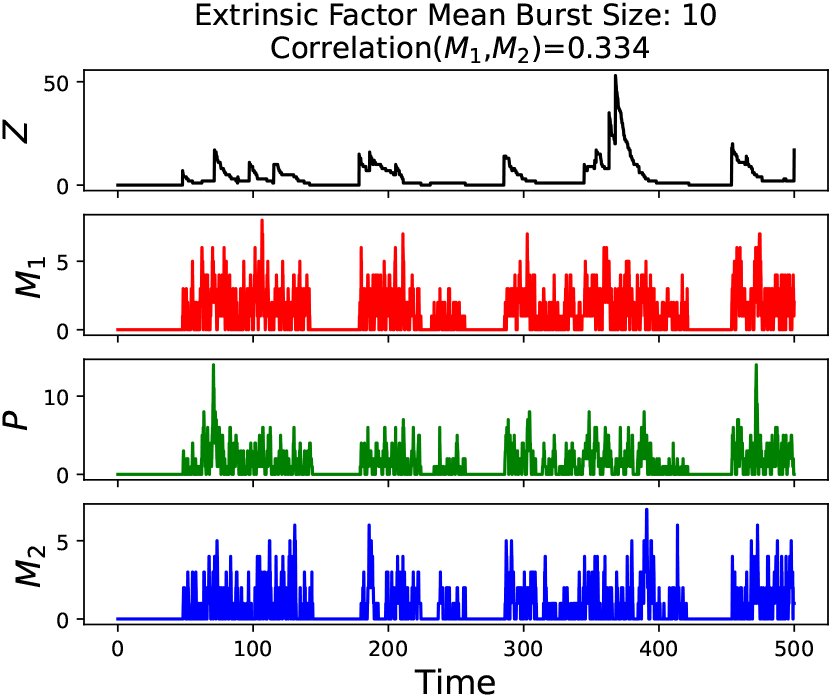
Simulation using parameter values listed in Table IV, with Extrinsic Factor burst size *β* drawn from a geometric distribution with mean 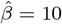. This simulation yielded a correlation coefficient between *M*_1_ and *M*_2_ of 0.334.

**Fig. 6.**
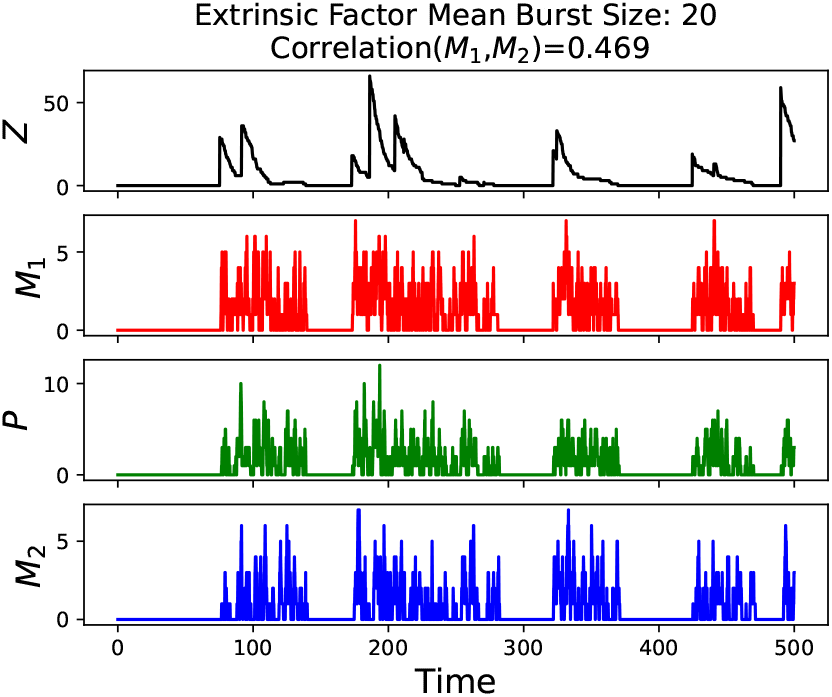
Simulation using parameter values listed in Table IV, with Extrinsic Factor burst size *β* drawn from a geometric distribution with mean 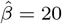.<This simulation yielded a correlation coefficient between *M*_1_ and *M*_2_ of 0.469.

Figure 7 shows the general relationship between the mean burst size 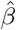 and the correlation between *M*_1_ and *M*_2_, and confirms what we could see visually in Figures 4, 5, and 6: higher mean burst size 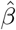 leads to higher levels of correlation between *M*_1_ and *M*_2_. Figure 8 shows this correlation for both different mean burst sizes of 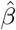 and different Protein/mRNA1 stability ratios (*τ*). Here, correlation levels for the Cascade scenario are compared to the Activation scenario results from Figure 2. Under conditions of extrinsic noise with high bursts of Extrinsic Factor production, the level of correlation between *M*_1_ and *M*_2_ persists more than in the scenario with no extrinsic noise, although it becomes slightly weaker.

**Fig. 7.**
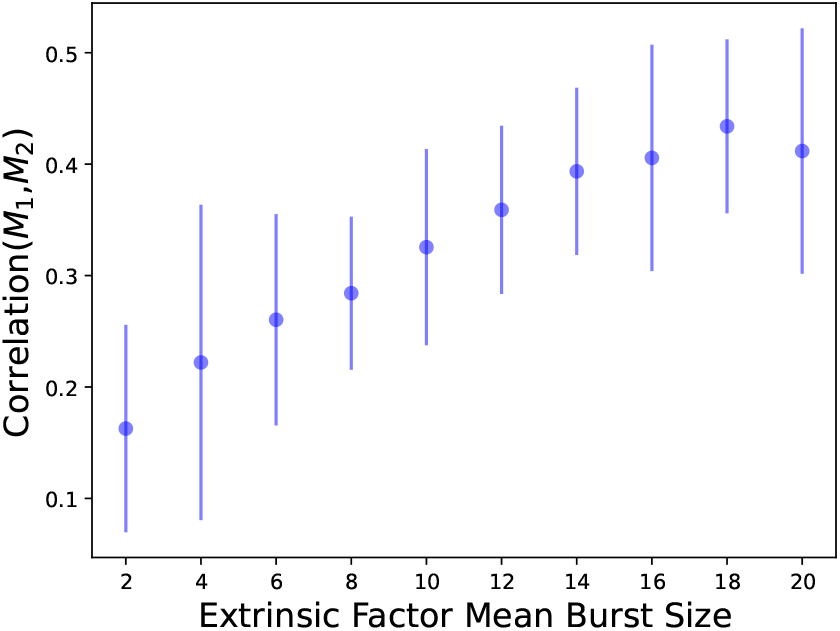
Simulations using parameter values listed in Table IV, with *β* drawn from geometric distributions with means ranging from 2 to 20 (as shown on the horizontal axis). The stochastic model for the Cascade scenario is described in Table III. Error bars show one standard deviation.

**Fig. 8.**
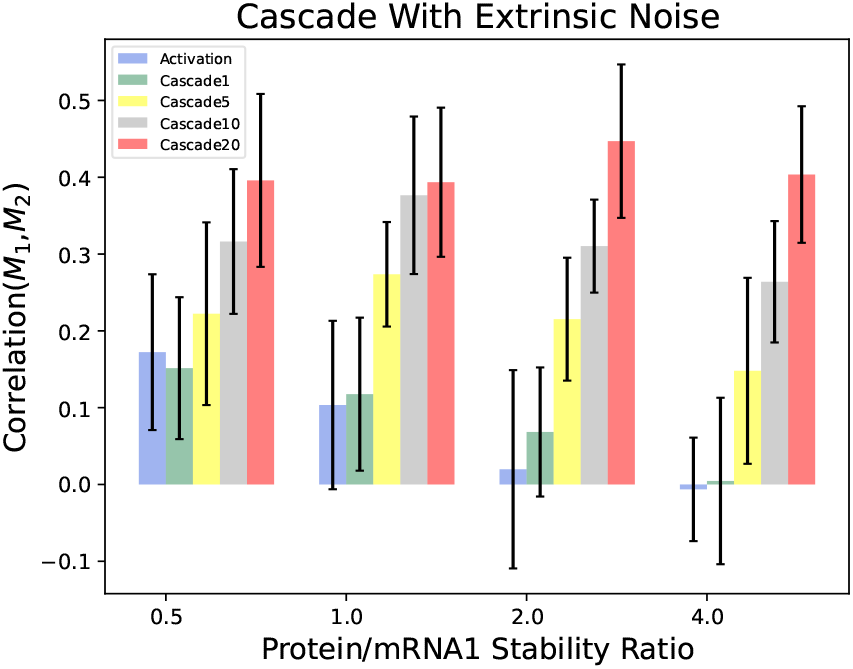
Simulations using parameter values listed in Table IV, with *β* drawn from geometric distributions of 1, 5, 10, and 20, and stability ratios (*τ*) of 0.5, 1, 2, and 4. Error bars show one standard deviation. The “Activation” bars show correlation levels for the Activation scenario with no extrinsic noise (shown previously in Figure 2). The bars labeled “Cascade1,” “Cascade5,” “Cascade10,” and “Cascade20” show the correlation levels for the Cascade scenario, with mean Extrinsic Factor burst sizes of 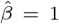, 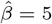 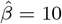, and 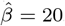, respectively.

In this section, we purposely defined the Extrinsic Factor in abstract terms without a definite biological meaning. However, it is interesting to consider possible biological implications of these results. Note that as we increase the burst size of Extrinsic Factor production, the model begins to resemble a system with distinct transcriptional states, rather than stochastic fluctuations around a single steady state, as in the Activation scenario with only intrinsic noise. For example, in Figure 6, the system could be thought of as representing the switching between two expression states (an ON state and an OFF state in this case). This phenomenon of transient switching between distinct gene expression states is thought to play a role in many biological systems, including drug resistance in cancer [30], [31], so it is interesting to note that GRN inference from mRNA abundance data may be more feasible under these conditions than under a single steady state condition.

## V. Distinguishing Between Cascade and Coregulation

Another key finding from Mahajan et al. 2022 [21] related to the difficulty of distinguishing between scenarios in which one gene regulates another and scenarios in which both genes are regulated by a common upstream regulator. Both of these scenarios can yield a correlation between the mRNA levels, so there is a possibility of a false positive network inference error in the latter scenario. In this section, we analyze a situation in which rather than Gene 1 regulating Gene 2, instead Gene 1 and Gene 2 are both regulated by the Extrinsic Factor. We will attempt to distinguish between this scenario and the previous scenarios in which Gene 1 directly regulated Gene 2.

In this new model, we continue using the Extrinsic Factor (tracked by *Z*) to model extrinsic noise, which can be thought of as an upstream regulator in this case. However, instead of the Extrinsic Factor activating Gene 1, which then activates Gene 2, in this model the Extrinsic Factor activates both Gene 1 and Gene 2 directly, with no direct regulation between Gene 1 and Gene 2. We refer to this as the Coregulation scenario. A diagram of this scenario is shown in Figure 9.

**Fig. 9.**
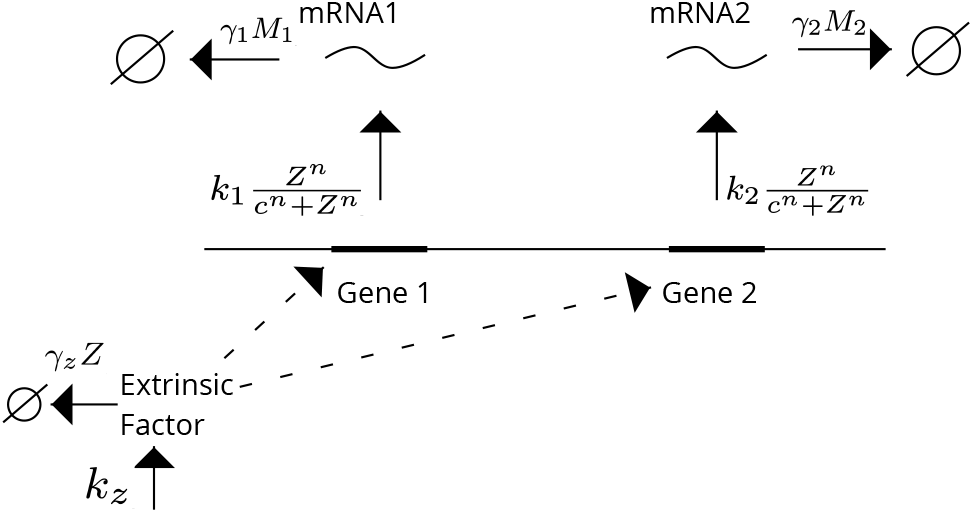
Coregulation Model. We can think of the Extrinsic Factor (tracked by *Z*) as an upstream regulator in this case. The Extrinsic Factor activates the transcription of mRNA1 (tracked by *M*_1_) and mRNA2 (tracked by *M*_2_), both of which also degrade with some rate. In this model, we no longer track the level of the Protein, because we do not need this information to model the production of mRNA2.

The stochastic model is described in Table V. We model the production and degradation of the Extrinsic Factor the same as in the last section, with production occurring in bursts of size *β*, drawn from a geometric distribution with mean 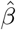. As in the last section, the transcription of both mRNA1 and mRNA2 is modeled with Hill functions, and the Hill function parameters *c* and *n* are the same for both. However, unlike in the previous section, the transcription of mRNA2 is now activated by the Extrinsic Factor, not by the Protein. The Protein is left out of this model since we no longer need to track its abundance to model the transcription of mRNA2.

**TABLE V.** Coregulation Model

| Event | Count Update | Propensity |
| --- | --- | --- |
| Extrinsic Factor production | $Z \rightarrow Z + \beta$ | $k_z$ |
| Extrinsic Factor degradation | $Z \rightarrow Z - 1$ | $\gamma_z Z$ |
| mRNA1 production | $M_1 \rightarrow M_1 + 1$ | $k_1 \frac{Z^n}{c^n + Z^n}$ |
| mRNA1 degradation | $M_1 \rightarrow M_1 - 1$ | $\gamma_1 M_1$ |
| mRNA2 production | $M_2 \rightarrow M_2 + 1$ | $k_2 \frac{Z^n}{c^n + Z^n}$ |
| mRNA2 degradation | $M_2 \rightarrow M_2 - 1$ | $\gamma_2 M_2$ |

Figure 10 shows the correlation between *M*_1_ and *M*_2_ for different burst sizes in the Coregulation scenario, compared to the correlation in the Cascade scenario that we simulated in the previous section. As you can see, it is very difficult to distinguish between the Cascade scenario and the Coregulation scenario based on only the correlation between the two mRNA levels.

**Fig. 10.**
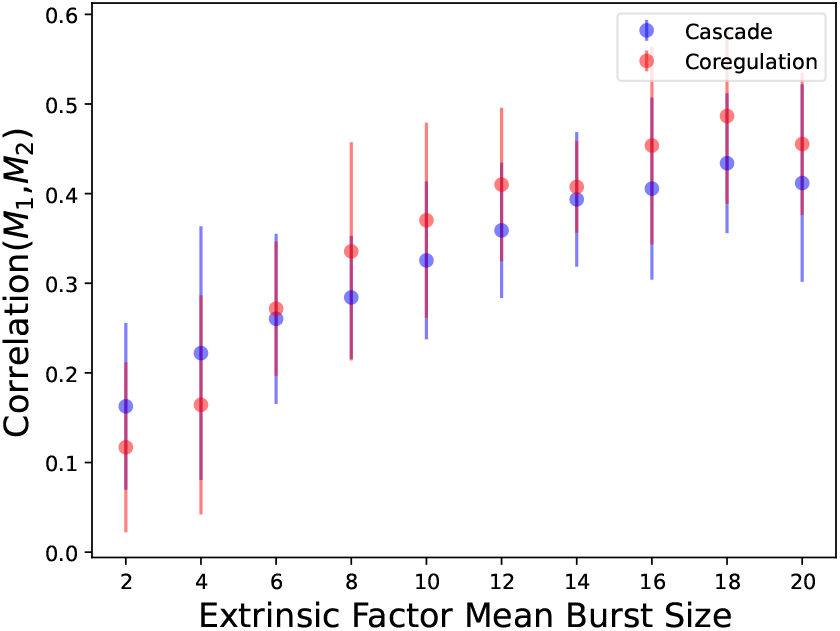
Simulations using parameter values listed in Table IV, with *β* drawn from geometric distributions with means ranging from 2 to 20 (as shown on the horizontal axis). The stochastic model for the Cascade scenario is described in Table III. The stochastic model for the Coregulation scenario is described in Table V. Blue dots show the correlation between *M*_1_ and *M*_2_ levels for the Cascade scenario, and red dots show the correlation for the Coregulation scenario. Error bars show one standard deviation.

However, it may be possible to distinguish between these scenarios based on the noise profiles of *M*_1_ and *M*_2_. Figure 11 shows the ratio 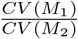 in both scenarios, for burst sizes ranging from 2 to 20. *CV* here is the coefficient of variation, or the standard deviation of the sample divided by the mean of the sample. It appears that we can distinguish between the scenarios using this noise ratio, even though both scenarios have similar levels of correlation. In the Coregulation scenario, *M*_1_ and *M*_2_ have similar noise, so their CV ratio is close to 1. However, in the Cascade scenario, *M*_1_ has lower noise than *M*_2_, leading to a lower CV ratio.

**Fig. 11.**
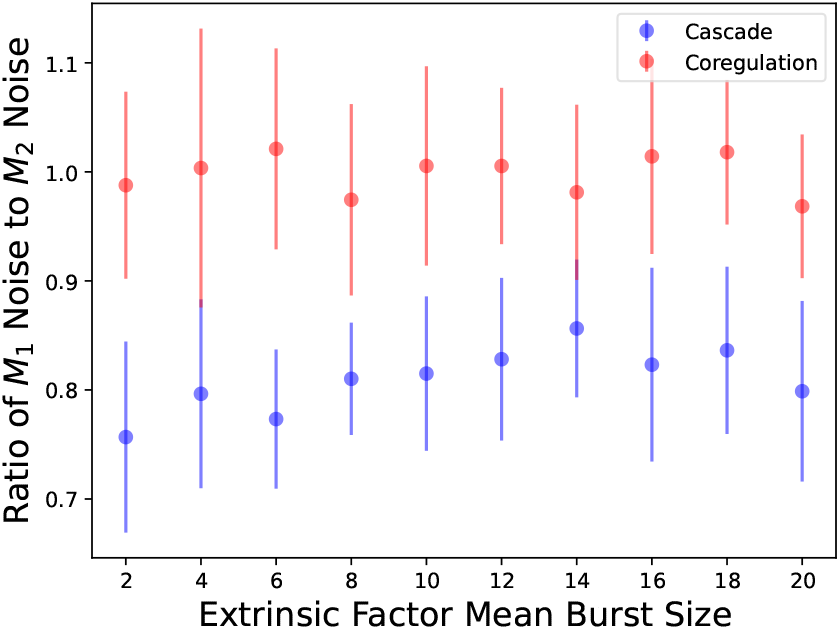
Simulations using parameter values listed in Table IV, with *β* drawn from geometric distributions with means ranging from 2 to 20 (as shown on the horizontal axis). In this case, rather than plotting the correlation between *M*_1_ and *M*_2_, we plot the noise ratio 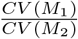, where *CV* is the coefficient of variation. Blue dots show the noise ratio for the Cascade scenario, and red dots show the noise ratio for the Coregulation scenario. Error bars show one standard deviation.

## VI. Further Analysis

In the previous sections, we used stochastic simulations of different scenarios to study the correlation between *M*_1_ and *M*_2_ under different noise conditions. While stochastic simulations are a valuable tool for analysis, it can also be helpful to have a mathematical framework for analysis that does not rely on simulations. In this section, we analyze simplified linear models of the three previously described scenarios, and derive formulas for the coefficients of variation of *M*_1_ and *M*_2_, and the correlation between *M*_1_ and *M*_2_ for each scenario.

### Activation

We begin with the first scenario in which Gene 1 regulates Gene 2, with no extrinsic noise. In our previous model for this scenario, we used a nonlinear Hill function to describe the activation of Gene 2 by Gene 1. However, in this section, we will make the simplifying assumption of a linear regulatory relationship between Gene 1 and Gene 2, in order to make the model more amenable to mathematical analysis. We also make the simplifying assumption that mRNA1, Protein, and mRNA2 all have the same degradation rate, which we call *γ*. After making these assumptions, the stochastic model for this scenario is described in Table VI.

**TABLE VI.** Linear Activation Model

| Event | Count Update | Propensity |
| --- | --- | --- |
| mRNA1 production | $M_1 \rightarrow M_1 + 1$ | $k_1$ |
| mRNA1 degradation | $M_1 \rightarrow M_1 - 1$ | $\gamma M_1$ |
| Protein production | $P \rightarrow P + 1$ | $k_p M_1$ |
| Protein degradation | $P \rightarrow P - 1$ | $\gamma P$ |
| mRNA2 production | $M_2 \rightarrow M_2 + 1$ | $k_2 P$ |
| mRNA2 degradation | $M_2 \rightarrow M_2 - 1$ | $\gamma M_2$ |

Our eventual goal is to derive a formula for the correlation between *M*_1_ and *M*_2_ in this model, as well as formulas for the coefficients of variation for both *M*_1_ and *M*_2_. In order to do this, we start by deriving the first and second order steady state moments for all of the variables. For the rest of this section, we will use angle brackets to signify expected value. For example, *M*_1_ will denote the expected value of 〈*M*_1_〉 (also called the first order moment of *M*_1_), and 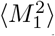 will denote the expected value of 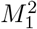 (also called the second order moment of *M*_1_).

With this simplified linear Activation model, we can use standard moment analysis techniques [10], [12], [33]–[35], [42] to solve for the first and second order steady state moments of *M*_1_ and *M*_2_:

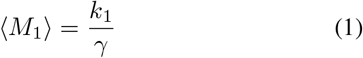

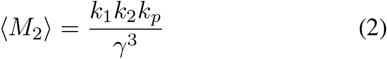

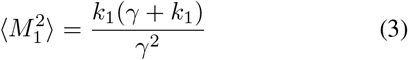

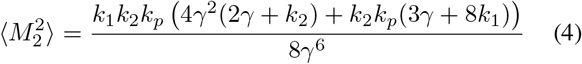

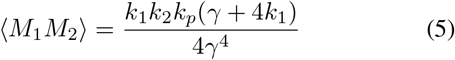

The coefficient of variation for a random variable can be written in terms of its first and second and order moments.

For example, the coefficient of variation for *M*_1_ can be written as:

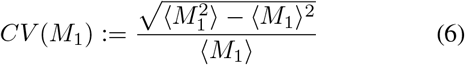

Since we already have expressions for the first and second order moments of *M*_1_ and *M*_2_, we can write the coefficients of variation for these variables in terms of the model parameters:

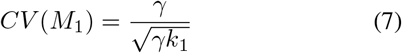

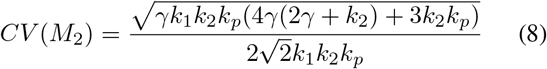

We note that, based on Equation 7, the variation in *M*_1_ is Poissonian, since 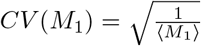 We can also write the correlation between *M*_1_ *M*_2_ in terms of the first and second order moments:

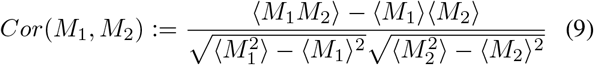

We can write this in terms of the model parameters:

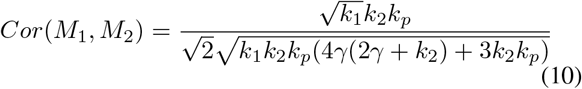

### Cascade

We use the same approach to derive formulas for these measures in the Cascade model, in which the Extrinsic Factor activates Gene 1, which then activates Gene 2. Again, we make the simplifying assumptions of linear activation rather than nonlinear activation via a Hill function, and that the mRNA1, Protein, and mRNA2 all have the same degradation rate, which we call *γ*. Also, we no longer model Extrinsic Factor production as a burst, so the *Z* production count update is now *Z → Z* + 1, not *Z → Z* + *β*. After making these assumptions, the stochastic model for the Cascade scenario is shown in Table VII.

**TABLE VII.** Linear Cascade Model

| Event | Count Update | Propensity |
| --- | --- | --- |
| Extrinsic Factor production | $Z \rightarrow Z + 1$ | $k_z$ |
| Extrinsic Factor degradation | $Z \rightarrow Z - 1$ | $\gamma_z Z$ |
| mRNA1 production | $M_1 \rightarrow M_1 + 1$ | $k_1 Z$ |
| mRNA1 degradation | $M_1 \rightarrow M_1 - 1$ | $\gamma M_1$ |
| Protein production | $P \rightarrow P + 1$ | $k_p M_1$ |
| Protein degradation | $P \rightarrow P - 1$ | $\gamma P$ |
| mRNA2 production | $M_2 \rightarrow M_2 + 1$ | $k_2 P$ |
| mRNA2 degradation | $M_2 \rightarrow M_2 - 1$ | $\gamma M_2$ |

We can use the same moment analysis techniques described in the previous section to write formulas for the coefficients of variation and correlation in the Cascade scenario. The coefficient of variation for *M*_1_ is:

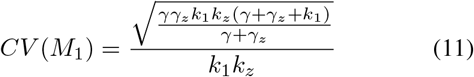

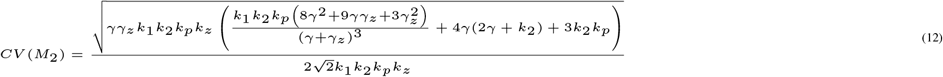

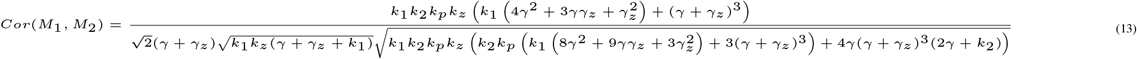

The coefficient of variation for *M*_2_ and the correlation between *M*_1_ and *M*_2_ are shown at the top of the next page because of their large size.

### Coregulation

Once again, we use the same approach for the Coregulation scenario, in which the Extrinsic Factor activates both Gene 1 and Gene 2. We again make the simplifying assumptions that this is linear activation, rather than activation via Hill function, and that the mRNA1, Protein, and mRNA2 all have the same degradation rate, which we call *γ*. Again, we no longer model *Z* production as a burst, so the *Z* production count update is now *Z → Z* +1, not *Z → Z* + *β*. After making these assumptions, the stochastic model for the Coregulation scenario is shown in Table VIII.

**TABLE VIII.** Linear Coregulation Model

| Event | Count Update | Propensity |
| --- | --- | --- |
| Extrinsic Factor production | $Z \rightarrow Z + 1$ | $k_z$ |
| Extrinsic Factor degradation | $Z \rightarrow Z - 1$ | $\gamma_z Z$ |
| mRNA1 production | $M_1 \rightarrow M_1 + 1$ | $k_1 Z$ |
| mRNA1 degradation | $M_1 \rightarrow M_1 - 1$ | $\gamma M_1$ |
| mRNA2 production | $M_2 \rightarrow M_2 + 1$ | $k_2 Z$ |
| mRNA2 degradation | $M_2 \rightarrow M_2 - 1$ | $\gamma M_2$ |

We use the same moment analysis techniques as in the previous sections to write expressions for the coefficients of variation and correlation for *M*_1_ and *M*_2_:

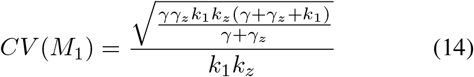

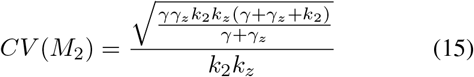

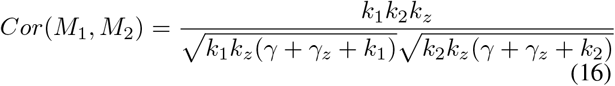

### Analytical Results

For this part of the analysis, we set the parameters so that 〈*M*_1_〉 = 〈*M*_2_〉, and so that this mean mRNA level is the same across all three scenarios. We then vary the mean mRNA level, and observe how the correlation and coefficients of variation change. In Figure 12, we show that as the mean mRNA level increases, the correlation between *M*_1_ and *M*_2_ increases in the Cascade and Coregulation scenarios, but decreases in the Activation scenario. In Figure 13, we show that as the mean mRNA expression level increases, the ratio of *M*_1_ noise to *M*_2_ noise holds steady at 1 in the Coregulation scenario, drops only very slightly before stabilizing in the Cascade scenario, and drops off quite steeply in the Activation scenario.

**Fig. 12.**
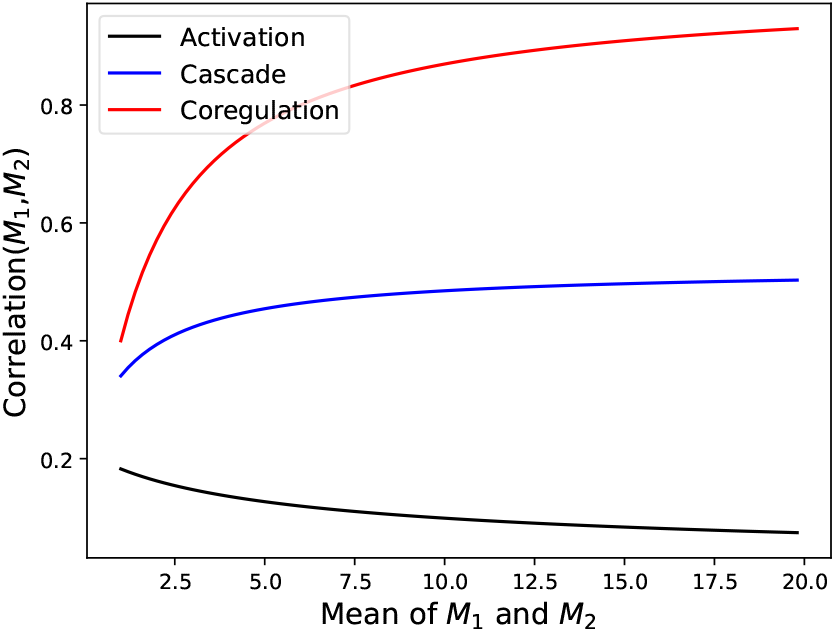
Correlation formula results for all three scenarios. We keep the mean mRNA abundance levels equal between *M*_1_ and *M*_2_, and equal across all three scenarios, and plot the correlation as we vary this mean mRNA expression level. Activation refers to the regulatory scenario described in Figure 1. Cascade refers to the regulatory scenario described in Figure 3. Coregulation refers to the regulatory scenario described in Figure 9.

**Fig. 13.**
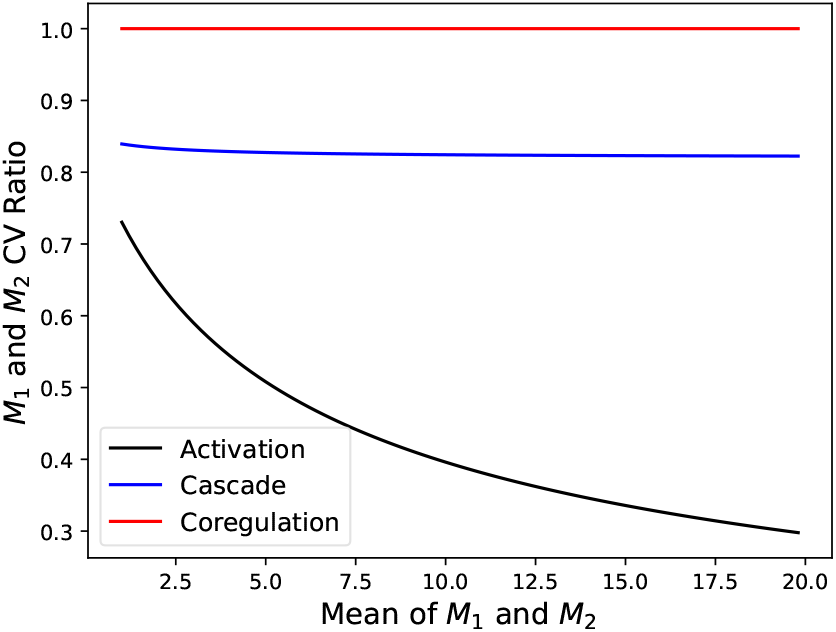
We keep the mean mRNA abundance levels equal between *M*_1_ and *M*_2_, and equal across all three scenarios, and plot the noise ratio 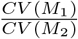 as we vary this mean mRNA expression level. Activation refers to the regulatory scenario described in Figure 1. Cascade refers to the regulatory scenario described in Figure 3. Coregulation refers to the regulatory scenario described in Figure 9.

These results have some interesting biological implications. They seem to suggest that the difference in feasibility of network inference between the Activation and Cascade scenarios is more pronounced in high copy number situations compared to low copy number situations, and that there is also more of a false positive threat from Coregulation scenario in high copy number situations. They also suggest that the differences in noise profiles between the direct regulation scenarios (Activation and Cascade) and the Coregulation scenario persist in high copy number situations.

## VII. Discussion

In this paper, we have analyzed the feasibility of network inference under different noise conditions through stochastic simulations, considering both intrinsic and extrinsic noise. We began by replicating a key result from Mahajan et al. 2022 [21] which suggests that under conditions of only intrinsic noise, the correlation between mRNA abundance levels for two genes in an activation relationship is quite weak, and gets weaker as the ratio of protein stability to mRNA stability increases. Under these conditions of only intrinsic noise, network inference from transcriptomic data would be very difficult.

Next, we investigated a scenario in which an extrinsic noise source activates the expression of a gene, which then activates the expression of another gene. Under these conditions, we found that the correlation between mRNA abundance levels for the two genes gets stronger as the extrinsic noise begins to resemble a state variable. We also found that the correlation persists (although it becomes weaker) even as the ratio of protein stability to mRNA stability increases. A biological takeaway from this result is that if a cell has distinct transient expression states, resulting from external factors or internal regulatory network dynamics, then under those conditions the task of network inference from transcriptomic data seems more tractable than under conditions of intrinsic noise only. This result is notable because transient state-switching between different gene expression states is thought to play a role in various biological phenomena, including drug resistance in cancer [30], [31].

We then considered a scenario in which two genes are coregulated by a common upstream gene. Under these conditions, we still observe a correlation between the mRNA abundance levels of the two genes, potentially leading to a false positive error in the task of network inference. However, simulation results suggest that even though this scenario yields similar levels of correlation to the Cascade scenario, we may be able to distinguish between the two scenarios using the noise profiles of the mRNA levels.

Finally, we provided a mathematical framework for further analysis of simplified linear models of each of the noise scenarios. We used moment analysis techniques to derive expressions for the coefficients of variation of the mRNA levels, as well as the correlation between them, and explored how these measures change with changes in mean mRNA levels. This allows us to make predictions about the difference between the feasibility of network inference in low copy number and high copy number situations, for each of the three noise scenarios.

Future work will include further theoretical analysis of these models. Additionally, goals for future work include testing the predictions made in this paper on real biological datasets, using data from single cell RNA-seq experiments, as well as further benchmarking of network inference methods on data model organisms and synthetic gene regulatory circuits.

## IX. Author Contributions Statement

M.S. and A.S. contributed to the analysis, writing, and editing of this paper.

## X. Acknowledgments

This work is supported by grants from the Army Research Office (W911NF1910243) and the National Science Foundation (ECCS-1711548).

## References

[1] M. Bansal, G. D. Gatta, and D. di Bernardo. Inference of gene regulatory networks and compound mode of action from time course gene expression profiles. Bioinformatics, 22(7):815–822, 2006.

[2] Shohag Barman and Yung-Keun Kwon. A Boolean network inference from time-series gene expression data using a genetic algorithm. Bioinformatics, 34(17):i927–i933, 2018.

[3] Ceélia Biane, Franck Delaplace, and Tarek Melliti. Abductive network action inference for targeted therapy discovery. Electronic Notes in Theoretical Computer Science, 335:3–25, 2018.

[4] A. J. Butte and I. S. Kohane. Mutual information relevance networks: Functional genomic clustering using pairwise entropy measurements. Biocomputing 2000, 1999.

[5] Thalia E. Chan, Michael P.H. Stumpf, and Ann C. Babtie. Gene regulatory network inference from single-cell data using multivariate information measures. Cell Systems, 5(3), 2017.

[6] Michael B. Elowitz, Arnold J. Levine, Eric D. Siggia, and Peter S. Swain. Stochastic gene expression in a single cell. Science, 297(5584):1183–1186, 2002.

[7] Jeremiah J Faith, Boris Hayete, Joshua T Thaden, Ilaria Mogno, Jamey Wierzbowski, Guillaume Cottarel, Simon Kasif, James J Collins, and Timothy S Gardner. Large-scale mapping and validation of Escherichia coli transcriptional regulation from a compendium of expression profiles. PLoS Biology, 5(1), 2007.

[8] Nir Friedman, Michal Linial, Iftach Nachman, and Dana Pe’er. Using bayesian networks to analyze expression data. Journal of Computational Biology, 7(3-4):601–620, 2000.

[9] Daniel T Gillespie. A general method for numerically simulating the stochastic time evolution of coupled chemical reactions. Journal of Computational Physics, 22(4):403–434, 1976.

[10] Daniel T. Gillespie. Approximate accelerated stochastic simulation of chemically reacting systems. The Journal of Chemical Physics, 115(4):1716–1733, 2001.

[11] Anne-Claire Haury, Fantine Mordelet, Paola Vera-Licona, and Jean-Philippe Vert. Tigress: Trustful inference of gene regulation using stability selection. BMC Systems Biology, 6(1), 2012.

[12] João Pedro Hespanha and Abhyudai Singh. Stochastic models for chemically reacting systems using polynomial stochastic hybrid systems. International Journal of Robust and Nonlinear Control, 15(15):669–689, 2005.

[13] Andreas Hilfinger and Johan Paulsson. Separating intrinsic from extrinsic fluctuations in dynamic biological systems. Proceedings of the National Academy of Sciences, 108(29):12167–12172, 2011.

[14] Vân Anh Huynh-Thu, Alexandre Irrthum, Louis Wehenkel, and Pierre Geurts. Inferring regulatory networks from expression data using tree-based methods. PLoS ONE, 5(9), 2010.

[15] Vân Anh Huynh-Thu and Guido Sanguinetti. Combining tree-based and dynamical systems for the inference of gene regulatory networks. Bioinformatics, 31(10):1614–1622, 2015.

[16] Vân Anh Huynh-Thu and Guido Sanguinetti. Gene regulatory network inference: An introductory survey. Methods in Molecular Biology, page 1–23, 2018.

[17] Guy Karlebach and Ron Shamir. Modelling and analysis of gene regulatory networks. Nature Reviews Molecular Cell Biology, 9(10):770–780, 2008.

[18] KC Kishan, Rui Li, Feng Cui, Qi Yu, and Anne R Haake. Gne: A deep learning framework for gene network inference by aggregating biological information. BMC Systems Biology, (13):38, 2019.

[19] P. K. Kreeger and D. A. Lauffenburger. Cancer systems biology: A network modeling perspective. Carcinogenesis, 31(1):2–8, 2009.

[20] Yansheng Liu, Andreas Beyer, and Ruedi Aebersold. On the dependency of cellular protein levels on mRNA abundance. Cell, 165(3):535–550, 2016.

[21] Tarun Mahajan, Michael Saint-Antoine, Roy D. Dar, and Abhyudai Singh. Limits on inferring gene regulatory networks from single-cell measurements of unstable mrna levels. 2022 IEEE 61st Conference on Decision and Control (CDC), 2022.

[22] Tarun Mahajan, Abhyudai Singh, and Roy D. Dar. Kinetic constraints on noise reduction in feedback gene regulatory networks. 2022 American Control Conference (ACC), 2022.

[23] Daniel Marbach, James C Costello, Robert Kuüffner, Nicole M Vega, Robert J Prill, Diogo M Camacho, Kyle R Allison, The DREAM5 Consortium, Manolis Kellis, James J Collins, and et al. Wisdom of crowds for robust gene network inference. Nature Methods, 9(8):796–804, 2012.

[24] Adam A Margolin, Ilya Nemenman, Katia Basso, Chris Wiggins, Gustavo Stolovitzky, Riccardo Dalla Favera, and Andrea Califano. ARACNE: An algorithm for the reconstruction of gene regulatory networks in a mammalian cellular context. BMC Bioinformatics, 7(S1), 2006.

[25] Patrick E. Meyer, Kevin Kontos, Frederic Lafitte, and Gianluca Bontempi. Information-theoretic inference of large transcriptional regulatory networks. EURASIP Journal on Bioinformatics and Systems Biology, 2007:1–9, 2007.

[26] Aditya Pratapa, Amogh P. Jalihal, Jeffrey N. Law, Aditya Bharadwaj, and T. M. Murali. Benchmarking algorithms for gene regulatory network inference from single-cell transcriptomic data. Nature Methods, 17(2):147–154, 2020.

[27] Mark Ptashne. The chemistry of regulation of genes and other things. Journal of Biological Chemistry, 289(9):5417–5435, 2014.

[28] Michael M. Saint-Antoine and Abhyudai Singh. Evaluating pruning methods in gene network inference. 2019 IEEE Conference on Computational Intelligence in Bioinformatics and Computational Biology (CIBCB), 2019.

[29] Michael M Saint-Antoine and Abhyudai Singh. Network inference in systems biology: Recent developments, challenges, and applications. Current Opinion in Biotechnology, 63:89–98, 2020.

[30] Lea Schuh, Michael Saint-Antoine, Eric M. Sanford, Benjamin L. Emert, Abhyudai Singh, Carsten Marr, Arjun Raj, and Yogesh Goyal. Gene networks with transcriptional bursting recapitulate rare transient coordinated high expression states in cancer. Cell Systems, 10(4), 2020.

[31] Sydney M. Shaffer, Margaret C. Dunagin, Stefan R. Torborg, Eduardo A. Torre, Benjamin Emert, Clemens Krepler, Marilda Beqiri, Katrin Sproesser, Patricia A. Brafford, Min Xiao, and et al. Rare cell variability and drug-induced reprogramming as a mode of cancer drug resistance. Nature, 546(7658):431–435, 2017.

[32] Vahid Shahrezaei, Julien F Ollivier, and Peter S Swain. Colored extrinsic fluctuations and stochastic gene expression. Molecular Systems Biology, 4(1):196, 2008.

[33] A. Singh and J.P. Hespanha. Models for multi-specie chemical reactions using polynomial stochastic hybrid systems. Proceedings of the 44th IEEE Conference on Decision and Control, 2005.

[34] Abhyudai Singh and João P. Hespanha. Stochastic hybrid systems for studying biochemical processes. Philosophical Transactions of the Royal Society A: Mathematical, Physical and Engineering Sciences, 368(1930):4995–5011, 2010.

[35] Abhyudai Singh and João P. Hespanha. Approximate moment dynamics for chemically reacting systems. IEEE Transactions on Automatic Control, 56(2):414–418, 2011.

[36] Abhyudai Singh and Mohammad Soltani. Quantifying intrinsic and extrinsic variability in stochastic gene expression models. PLoS ONE, 8(12), 2013.

[37] Nitin Singh and Mathukumalli Vidyasagar. Blars: An algorithm to infer gene regulatory networks. IEEE/ACM Transactions on Computational Biology and Bioinformatics, 13(2):301–314, 2016.

[38] Peter S. Swain, Michael B. Elowitz, and Eric D. Siggia. Intrinsic and extrinsic contributions to stochasticity in gene expression. Proceedings of the National Academy of Sciences, 99(20):12795–12800, 2002.

[39] Zahra Vahdat, Khem Raj Ghusinga, and Abhyudai Singh. Comparing feedback strategies for minimizing noise in gene expression event timing. 2021 29th Mediterranean Conference on Control and Automation (MED), 2021.

[40] Zahra Vahdat, Karol Nienaltowski, Zia Farooq, Michal Komorowski, and Abhyudai Singh. Information processing in unregulated and autoregulated gene expression. 2020 European Control Conference (ECC), 2020.

[41] Yue Wang and Zikun Wang. Inference on the structure of gene regulatory networks. Journal of Theoretical Biology, 539:111055, 2022.

[42] Darren J. Wilkinson. Stochastic modelling for systems biology. Taylor and Francis, 2012.

[43] Ting Xu, Le Ou-Yang, Xiaohua Hu, and Xiao-Fei Zhang. Identifying gene network rewiring by integrating gene expression and gene network data. IEEE/ACM Transactions on Computational Biology and Bioinformatics, 15(6):2079–2085, 2018.

[44] J. Yu, V. A. Smith, P. P. Wang, A. J. Hartemink, and E. D. Jarvis. Advances to Bayesian network inference for generating causal networks from observational biological data. Bioinformatics, 20(18):3594–3603, 2004.

[45] Bin Zhang and Steve Horvath. A general framework for weighted gene co-expression network analysis. Statistical Applications in Genetics and Molecular Biology, 4(1), 2005.

[46] Haitao Zhao and Zhong-Hui Duan. Cancer genetic network inference using gaussian graphical models. Bioinformatics and Biology Insights, 13, 2019.

